# Vaginal microbiota of adolescents and their mothers: A preliminary study of vertical transmission and persistence

**DOI:** 10.1101/768598

**Authors:** Christine M. Bassis, Kaylie A. Bullock, Daniel E. Sack, Katie Saund, Ali Pirani, Evan S. Snitkin, Veronica I. Alaniz, Elisabeth H. Quint, Vincent B. Young, Jason D. Bell

**Affiliations:** Division of Infectious Disease, Department of Internal Medicine; Department of Microbiology and Immunology; Department of Obstetrics and Gynecology, University of Michigan, Ann Arbor, MI 48109

**Keywords:** Vaginal microbiota, transmission, birth mode, 16S rRNA gene sequences, *Lactobacillus crispatus* genomics

## Abstract

**Background:** Factors that influence vaginal microbiota composition, including its source, are not well understood. To determine if vaginal microbiota transmission from mother to daughter at birth influences the human vaginal microbiota composition in adolescence, we investigated the relationship between the vaginal microbiota of 13 mother/daughter pairs and the daughter’s birth mode.

**Results:** Based on analysis of bacterial 16S rRNA gene sequences, the vaginal microbiotas of mother/daughter pairs were more similar to each other if the daughter was born by vaginal delivery rather than by C-section. Additionally, genome sequences from an important member of the vaginal microbiota, *Lactobacillus crispatus*, isolated from one mother/daughter pair in which the daughter was born by vaginal delivery, were highly similar.

**Conclusions:** Both community-level analysis and isolate genome sequence analysis are consistent with birth-mode dependent transmission and persistence of at least some members of the vaginal microbiota.

**Importance:** The composition of the human vaginal microbiota is related to many aspects of health from infection susceptibility to preterm birth. Our study provides evidence that transmission of vaginal bacteria from mother to daughter at birth may be an important factor influencing vaginal microbiota composition into adolescence.

## Background

The vaginal microbiota plays an important role in human health. The community structure of the vaginal microbiota is linked to infection susceptibility and preterm birth (1–6). The composition of the vaginal microbiota is distinct from other body sites and contains types of bacteria that seem specific to the vagina 7). For example, the vaginal microbiota is often dominated by specific types of *Lactobacillus*, most commonly *L. crispatus* and *L. iners* (8, 9). Vaginal *Lactobacillus* sp. are thought to maintain dominance and inhibit colonization of other microbes through lactic acid production (10, 11).

Despite strong evidence that the vaginal microbiota can have significant impacts on health, the factors that influence the composition of the vaginal microbiota are not well understood. It is not known how this vagina-specific community is maintained from generation to generation. One possibility is that at least some members of the vaginal microbiota are transmitted from mother to daughter at birth and maintained in daughters through adolescence.

In healthy babies, the first large, direct exposure to microbes occurs at birth. Birth mode has been shown to influence the composition of the newborn microbiota (gut, skin, mouth), likely due to different bacterial exposure in vaginal delivery and C-sections (12, 13). However, the effect of birth mode on the composition of the vaginal microbiota has not been investigated. In this study, we compared the vaginal microbiotas of 13 mother/daughter pairs and investigated the effect of birth mode on mother/daughter microbiota similarity. We also compared the genome sequences from *Lactobacillus crispatus* isolates from one mother/daughter pair. We hypothesized that the vaginal microbiota of mothers and daughters would be more similar if the daughter was born by vaginal delivery than by C-section.

## Methods

### Subject recruitment and sample collection

Mother/daughter pairs were recruited from the Pediatric and Adolescent Gynecology Clinic at the University of Michigan Health System in 2014 and 2015. Exclusions were pregnancy and age of less than 15 years. Written, informed consent was obtained and participants completed a baseline survey on their demographics and pertinent gynecologic and medical history. Vaginal samples were self-collected using a dual-headed swab (Starplex Scientific, S09D) at baseline and then weekly for 4 weeks. The baseline swab was obtained in the clinic, with immediate storage on ice and transfer to −80°C within a few hours. The subsequent swabs were returned via mail at ambient temperature. After the fifth swab was received and a completion incentive was mailed to the subject, the link between samples and subject names was destroyed, irreversibly de-identifying all samples. The study was approved by the University of Michigan IRB (HUM00086661).

### DNA isolation and 16S rRNA gene sequencing

One of the swab heads from each sample was clipped directly into the bead plate of a PowerMag Microbiome RNA/DNA Isolation Kit (Mo Bio Laboratories, Inc.). DNA isolation was performed according the manufacturer’s instructions using an epMotion 5075 liquid handling system. The V4 region of the 16S rRNA gene was amplified from 1 or 7μl DNA and sequenced with a MiSeq (Illumina, San Diego, CA) using the 500 cycle MiSeq Reagent Kit, v. 2 (Illumina, catalog No. MS-102-2003) by the University of Michigan Microbial Systems Molecular Biology Laboratory as described previously (14). The other swab head was used for cultivation or stored at −80°C.

### Bacterial community analysis

The 16S rRNA gene sequences were processed using mothur v.1.36.1 and v.1.39.5 following the mothur MiSeq SOP (15, 16). Details of the processing steps are available in mother.daughter mothur.batch (https://github.com/cbassis/MotherDaughter_Vaginal_Microbiota.study). After sequence processing and alignment to the SILVA reference alignment (Release 102) (17), sequences were binned into operational taxonomic units (OTUs) based on 97% sequence similarity using the average neighbor method (18, 19). Samples with fewer than 1000 sequences were excluded from the analysis. OTUs were classified to the genus level within mothur using a modified version of the Ribosomal Database Project (RDP) training set (version 9) (20, 21). To further classify the *Lactobacillus* OTUs, representative sequences were analyzed using standard nucleotide BLAST for highly similar sequences (megablast) on the National Center for Biotechnology Information (NCBI) BLAST web page (https://blast.ncbi.nlm.nih.gov/Blast.cgi) (22). OTU relative abundances were calculated and plotted in a heatmap. To compare bacterial communities between pairs, within pairs and within subjects, we calculated θ_YC_ distances (a metric that takes relative abundances of both shared and non-shared OTUs into account) (23). A Kruskal-Wallis test with a Dunn’s posttest or a Wilcoxon (Mann-Whitney) test were used to determine if differences in θ_YC_ distances were statistically significant. Principal coordinates analysis (PCoA) was used to visualize the θ_YC_ distances between samples. R Studio (Version 1.1.456) with R (Version 3.5.1) was used for the statistical tests and plotting the heat map, box and whisker plots, and the ordination using the code available: https://github.com/cbassis/MotherDaughter_Vaginal_Microbiota.study/tree/master/R_co_de. Adobe Illustrator (CS6) was used for labeling and formatting figures.

### *Lactobacillus crispatus* isolation

For pair I, the second swab head from the freshly collected baseline vaginal sample was swabbed onto an MRS agar plate and incubated in an anaerobic chamber (Coy Laboratory Products) at 37°C. Individual isolates were identified via Sanger sequencing of the near-full length 16S rRNA gene.

### DNA isolation and genome sequencing

Three *Lactobacillus crispatus* isolates from pair I, 2 from the mother and 1 from the daughter, were grown overnight in 1 ml liquid MRS in an anaerobic chamber (Coy Laboratory Products) at 37°C. Genomic DNA was isolated from the liquid cultures using the PowerMicrobiome™ RNA Isolation Kit (Mo Bio Laboratories, Inc.) without the DNase treatment. Genome sequencing was performed by the Microbial Systems Molecular Biology Laboratory at the University of Michigan using an Illumina Nextera™ sequencing kit and a MiSeq (Illumina, San Diego, CA).

### Genome sequence analysis

Phylogenetic relationships between *L. crispatus* isolates from mother/daughter pair I and all *L. crispatus* strains with genome sequences available as fastq files from NCBI on December 27th, 2018 were determined based on recombination-filtered single nucleotide polymorphisms (SNPs). Quality of reads was assessed with FastQC v0.11.3 (24), and Trimmomatic 0.36 (25) was used for trimming adapter sequences and low-quality bases. Variants were identified by (i) mapping filtered reads to reference genome sequence *L. crispatus* ST1 (SAMEA2272191) using the Burrows-Wheeler short-read aligner (bwa-0.7.17) (26, 27), (ii) discarding polymerase chain reaction duplicates with Picard (picard-tools-2.5.0) (28), and (iii) calling variants with SAMtools (samtools-1.2) and bcftools (29). Variants were filtered from raw results using GATK ‘s (GenomeAnalysisTK-3.3-0) VariantFiltration (QUAL, >100; MQ, >50; >=10 reads supporting variant; and FQ, <0.025) (30). In addition, a custom python script was used to filter out single-nucleotide variants that were (i) <5 base pairs (bp) in proximity to indels, (ii) fell under Phage and Repeat region of the reference genome (identified using Phaster (31) and Nucmer (MUMmer3.23) (32)), (iii) not present in the core genome, or (iv) in a recombinant region identified by Gubbins 2.3.1 (33). A maximum likelihood tree was constructed in RAxML 8.2.8 (34) using a general-time reversible model of sequence evolution. Bootstrap analysis was performed with the number of bootstrap replicates determined using the bootstrap convergence test and the autoMRE convergence criteria (-N autoMRE). Bootstrap support values were overlaid on the best scoring tree identified during rapid bootstrap analysis (-f a). The final maximum likelihood tree was plotted and pairwise SNP distances were calculated in R Studio (Version 1.1.463) with R (Version 3.5.3): https://github.com/cbassis/MotherDaughter_Vaginal_Microbiota.study/blob/master/R_co_de/Mother_Daughter_Figure_3_Genome_Tree_and_Genome_Analysis.Rmd. Adobe Illustrator (CS6) was used for labeling and formatting the figure.

### Calculation of doubling time estimate for vaginal *L. crispatus in vivo*

We used the number of SNPs between the pair I mother and daughter *L. crispatus* isolates to estimate the doubling time of vaginal *L. crispatus in vivo* if all SNPs in the recombination-filtered core genome were due to mutations acquired since the daughter’s birth: Doubling time=(mutation rate)(daughter’s age)(genome length)/(# of mutations) The mutation rate of *L. crispatus* is unknown, so for this estimate we used the published mutation rate of another *Lactobacillus, L. casei* Zhang, *in vitro*, without antibiotics (1.0×10^-9^ bp/generation) (35). The pair I daughter’s age in hours was: 175,200 hours =(20 years)(365 days/year)(24 hour/day). The average length of the recombination-filtered core genome (940,943 bp) was used for genome length. We assumed that the isolates arose from a common ancestor and that all mutations were non-convergent, so the number of mutations acquired by each isolate would equal the number of SNP_s_ between the mother’s isolate and the daughter’s isolate divided by 2. We also estimated the number of mutations acquired per isolate core genome per year as (# of mutations)/(daughter’s age)=(# of SNP_s_)/2(daughter’s age).

## Results

### Subject characteristics and sequencing results

A total of 107 self-collected, vaginal swab samples were obtained from 26 subjects (13 mother/daughter pairs) (Table 1). Each subject returned 1-5 weekly samples (median=5 samples/subject, IQR=1). After sequence processing and exclusion of samples with fewer than 1000 sequences, a total of 2,336,437 high quality bacterial 16S rRNA gene sequences from 101 samples were analyzed with an average of 23,133 +/-10,212 sequences per sample.

**Table 1.** Subject Characteristics.

| Table 1. Subject Characteristics |  |  |  |  |
| --- | --- | --- | --- | --- |
|  | Mother (n=13) |  | Daughter (n=13) |  |
|  | Daughter's birth mode: Vaginal (n=10) | Daughter's birth mode: C-section (n=3) | Daughter's birth mode: Vaginal (n=10) | Daughter's birth mode: C-section (n=3) |
| Age, mean ± SD, years | 44.8±5.6 | 54±2.4 | 17.1±2.0 | 18.7±1.9 |
| Race: White (vs. Black, Asian, Hispanic, other) | 90% (n=9) | 100% (n=3) | 90% (n=9) | 100% (n=3) |
| Subject Birth mode: Vaginal (vs. C-section) | 70% (n=7) | 100% (n=3) | 100% (n=10) | 0% (n=0) |
| Reproductive stage: Premenarchal | 0% (n=0) | 0% (n=0) | 10% (n=1) | 0% (n=0) |
| Reproductive stage: Reproductive | 70% (n=7) | 33% (n=1) | 90% (n=9) | 100% (n=3) |
| Reproductive stage: Postmenopausal | 30% (n=3) | 67% (n=2) | 0% (n=0) | 0% (n=0) |

### An individual’s vaginal microbiota is relatively stable over 4 weeks

During the sampling period, the vaginal microbiota of each subject was relatively stable. The high stability of the vaginal microbiota is apparent from the consistent within subject community composition (Figure 1). For example, high relative abundances of OTU1 (*L. crispatus*) and/or OTU2 (*L. iners*) persisted from week to week in many subjects. Additionally, average **θ**_YC_ distances were significantly lower within subjects than between subjects (Figure 2A) and samples clustered by subject in a PCoA based on **θ**_YC_ distances (Supplemental Figure 1).

**Figure 1.**
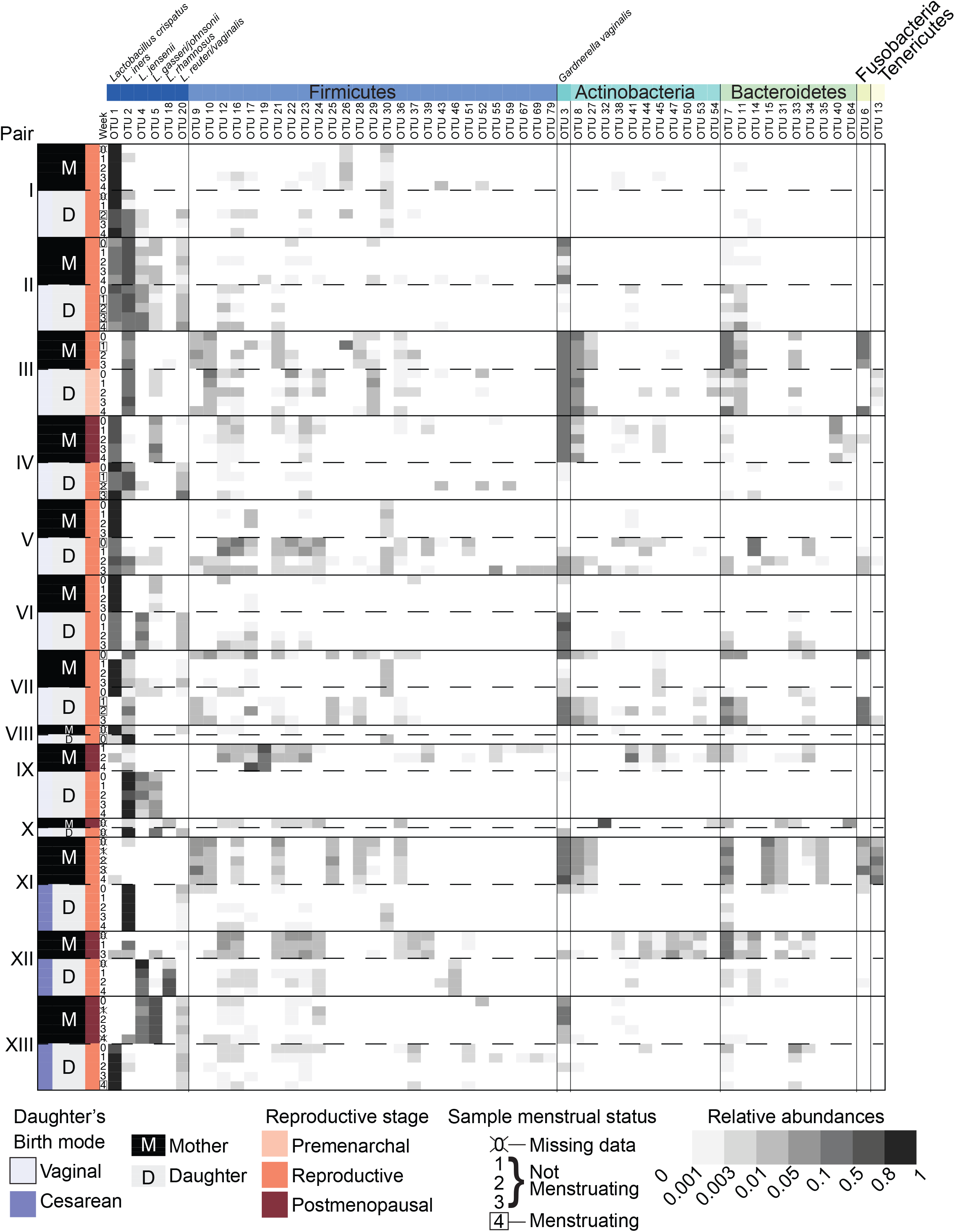
Vaginal bacterial community compositions of mother/daughter pairs. Relative abundances of OTUs in weekly vaginal swab samples from 13 mother/daughter pairs. Mother/daughter pairs were ordered by average within pair **θ**_YC_ distances, with the most similar pair (I) on top and the least similar pair (XIII) on the bottom. OTUs with a minimum of 200 sequences in the dataset overall and present at a relative abundance greater than 2% in at least 1 sample were included in the heat map.

**Figure 2.**
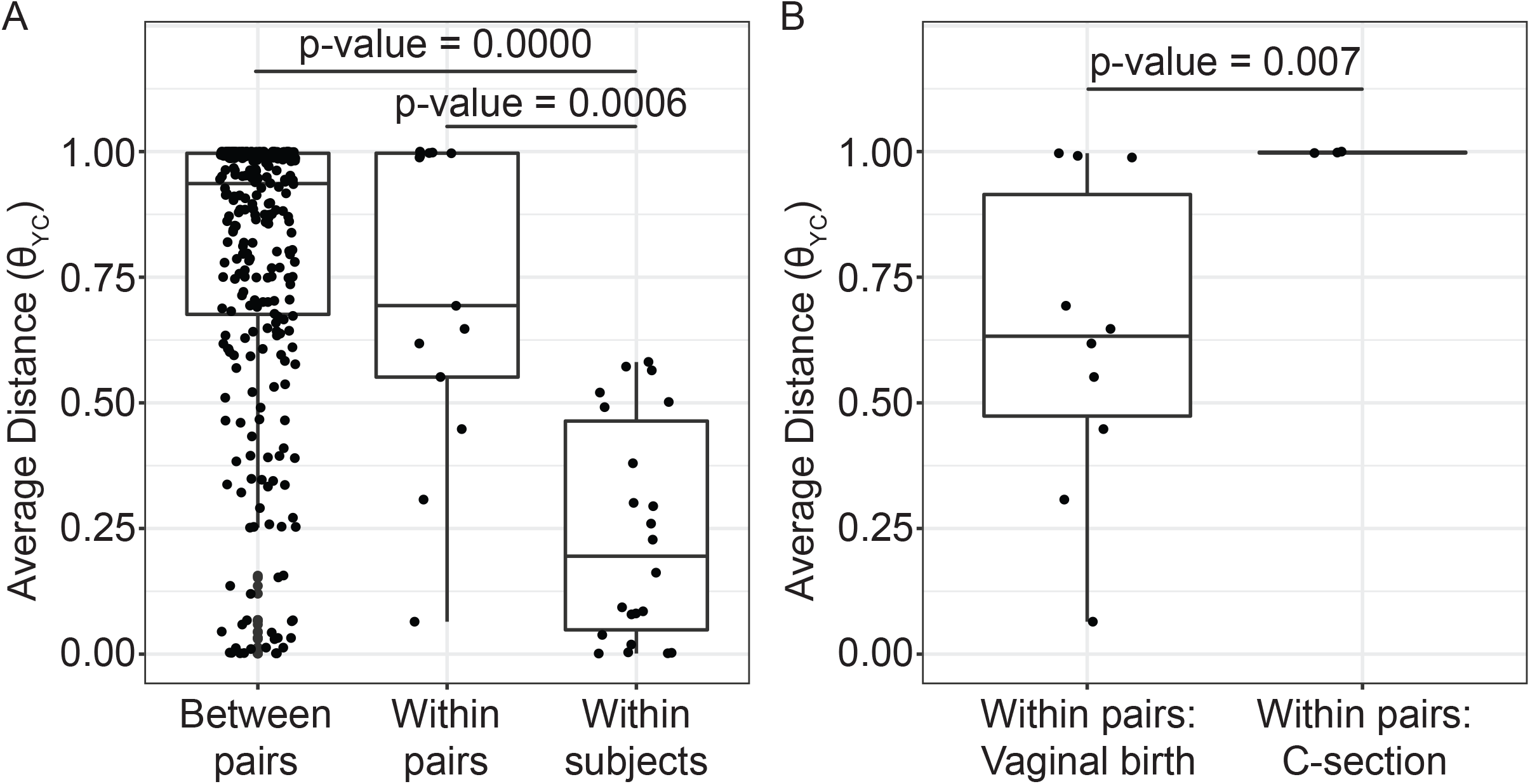
Average distances between vaginal bacterial communities. A. Average **θ**_YC_ distances between subjects from different mother/daughter pairs (between pairs), between subjects within a mother/daughter pair (within pair) and between samples from the same subject (within subject). P-values for comparisons that were significantly different by Dunn’s posttest are shown (Kruskal-Wallis p-value= 8.154e-10). B. Average **θ**_YC_ distances between subjects within a mother/daughter pair for daughters born by vaginal birth and by C-section. Wilcoxon (Mann-Whitney) test p-value is shown. In the box and whiskers plots, the median **θ**_YC_ distance is indicated by a line, values within the first to the third quartiles are inside the box and the whiskers extend to the smallest and largest values within 1.5x the interquartile range.

### Daughters born via vaginal delivery have greater microbiota similarity with their mothers than those born via C-section

To determine if mothers and their daughters had more similar vaginal microbiotas than unrelated subjects, we compared the average **θ**_YC_ distances between all unrelated subjects (between pairs) and the average **θ**_YC_ distances between mothers and their own daughters (within pairs) (Figure 2A). There was a trend toward greater similarity (lower **θ**_YC_ distances) within all mother/daughter pairs than between subjects in different mother/daughter pairs. To determine if birth mode was related to vaginal microbiota similarity within mother/daughter pairs, we compared the average within pair **θ**_YC_ distances for pairs in which the daughter was born by vaginal delivery and by C-section (Figure 2B). The average within pair **θ**_YC_ distances were significantly lower for pairs in which the daughter was born by vaginal delivery compared to C-section (Fig. 2B). Therefore, the vaginal microbiotas of daughters born by vaginal delivery were significantly more similar to their mothers’ than the daughters born by C-section were to their mothers’ (Fig. 2B).

### *Lactobacillus crispatus* isolates from mother/daughter pair I have highly similar genome sequences

The birth mode-dependent similarity of the vaginal microbiotas of mothers and their daughters suggested that vaginal bacteria could be transmitted between generations at birth and persist into adolescence. However, it is possible that genetic or environmental factors shared by a mother and her daughter lead to acquisition of similar bacteria later, resulting in the *de novo* establishment of similar vaginal communities. To investigate the possibility of direct transmission and persistence of one member of the vaginal microbiota, we generated draft genome sequences of *Lactobacillus crispatus* strains isolated from the freshly collected second swab head of mother/daughter pair I. The draft genome sequences of these isolates were compared with publicly available *L. crispatus* genome sequences by constructing a maximum likelihood phylogenetic tree based on a recombination-filtered core genome alignment. Interestingly, the three strains of *L. crispatus* from mother/daughter pair I, UMP1M1, UMP1M2 and UMP1D1, were more similar to each other than to any of the other strains, including others isolated from the female reproductive tract (Fig.3).

**Figure 3.**
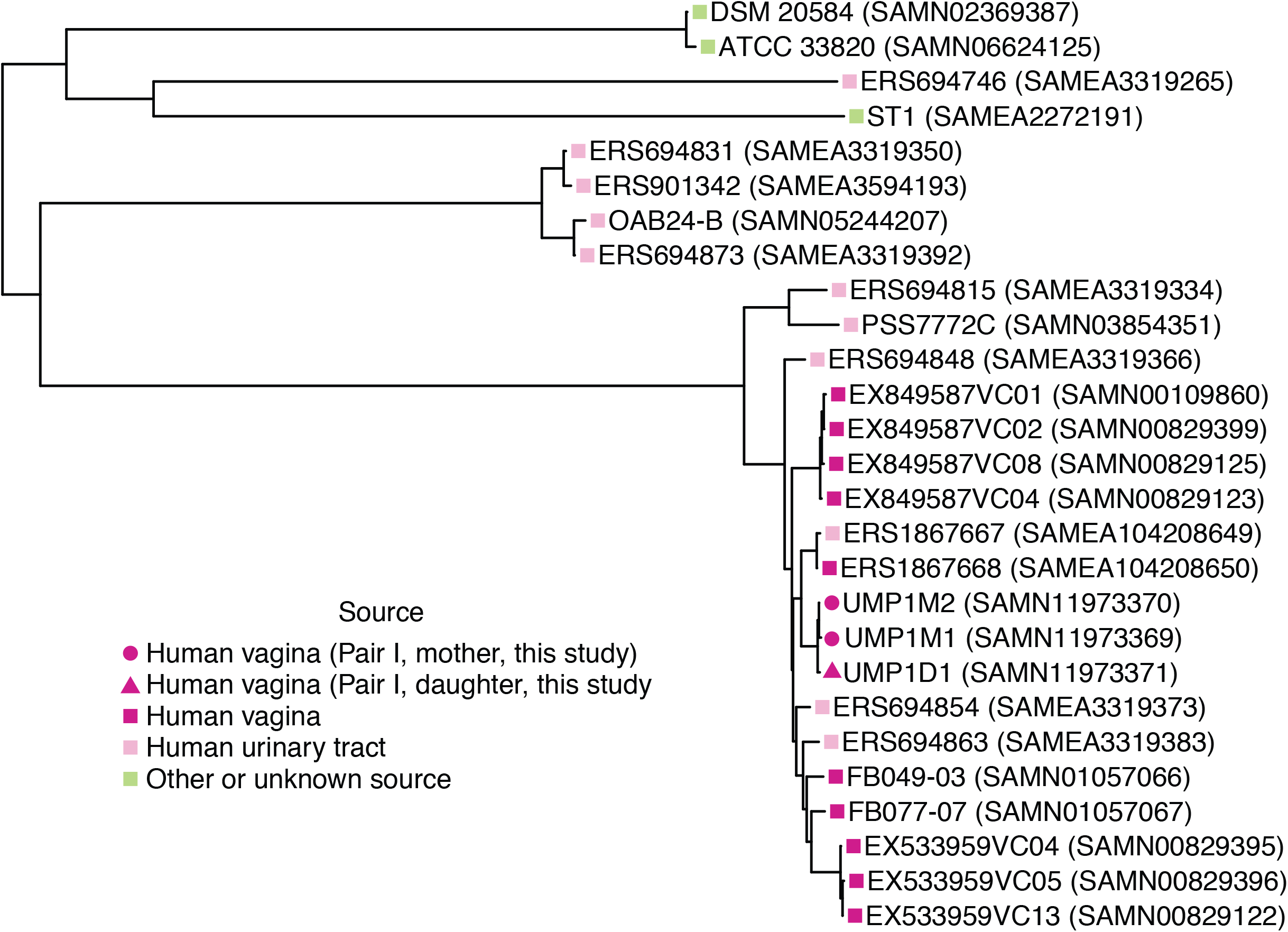
Phylogenetic relationships between *L. crispatus* strains. Maximum likelihood tree based on recombination-filtered SNP distances between *L. crispatus* genome sequences of isolates from mother/daughter pair I and other *L. crispatus* strains with publicly available genomes. Tip labels indicate *L. crispatus* strain names and NCBI BioSample identifiers. Bootstrap values were greater than or equal to 0.65.

We also calculated the number of SNPs between our isolates using the recombination-filtered core genome alignment. There were 11 recombination-filtered SNPs between the 2 isolates from the mother (UMP1M1 and UMP1M2) and 25 and 16 recombination-filtered SNPs between the daughter’s isolate (UMP1D1) and the 2 isolates from the mother (UMP1M1 and UMP1M2, respectively).

### Estimate of *in vivo* doubling time and mutation rate for vaginal *L. crispatus*

To further investigate the plausibility that the *L. crispatus* strain isolated from daughter I descended from a strain transmitted from her mother at birth, we estimated the doubling time that would allow our isolates to acquire the observed number of SNPs over 20 years. Based on the 25 SNPs between UMP1M1 and UMP1D1, the estimated doubling time for *L. crispatus in vivo* would be 13.2 hours. Based on the 16 SNPs between UMP1M2 and UMP1D1, the estimated doubling time would be 20.6 hours. We also estimated the *in vivo* mutation rate of the core genome of the *L. crispatus* isolates to be 0.4-0.6 mutations per year.

## Discussion

Our study provides preliminary evidence that the vaginal microbiota may be vertically transmitted from mother to daughter at birth via vaginal delivery and persists into adolescence. Because the daughters in our study were 15-21 years old, both transmission and persistence were required to observe evidence of vertical transmission. The first piece of evidence supporting vertical transmission is that the vaginal microbiotas of mothers and their adolescent daughters were more similar if their daughter was born by vaginal delivery rather than C-section. The second piece of evidence supporting vertical transmission and persistence is that an important member of the vaginal microbiota, *L. crispatus*, isolated from a vaginally-born, 20-year-old daughter and her mother (pair I) had highly similar genome sequences.

Other studies have compared the vaginal microbiotas of mothers and daughters without analyzing the effect of birth mode (36–38). One study found greater similarity between the vaginal microbiotas of mothers and daughters than between unrelated subjects (38). This was similar to the trend we observed toward greater community similarity within mother/daughter pairs, regardless of birth mode, than between unrelated subjects in different mother/daughter pairs (Figure 2A). Notable similarity between the vaginal microbiota of mothers and daughters was not detected in the other studies (36, 37). If many of the daughters in the other studies were born by C-section then high similarity between mothers and daughters would not be expected. With C-section rates of ~30% in the United States (study site for (37)) and ~36% in South Korea (study site for (36)) this is a possibility (39, 40). Additionally, our study focused on adolescent daughters (age 15-21) while the other studies focused on either younger or older daughters. Since reproductive stage seems to influence the structure of the vaginal microbiota (41), differences in reproductive stage may contribute to differences in vaginal community composition between mothers and daughters. Finally, we used a different method of comparing the vaginal microbiotas of mothers and daughters. We calculated distances between mothers and daughter using θ_YC_, a metric that accounts for the relative abundances of shared and non-shared OTUs, while the other studies were based on community types (37) and Unifrac (36). Although an overall community similarity was not observed in these studies, specific community members *(Lactobacillus* and *Prevotella*) were identified as most heritable in one study (36).

Based on the number of SNPs observed between the mother and daughter *L. crispatus* isolates and published mutations rates for *L. casei* Zhang (35), we estimated that *L. crispatus* would have an *in vivo* doubling time of 13.2-20.6 hours, depending on the specific isolates compared. The doubling time estimates of 13.2 hours and 20.6 hours for *L. crispatus in vivo* are within the range estimated for other bacteria in their natural environments, including *Escherichia coli* (15 hours) and *Salmonella enterica* (25 hours) (42). These doubling times are faster than the 4.1-5.6 days doubling times measured for *L. casei* Shirota in mouse intestines, where its growth rate was insufficient to maintain colonization (43). Although the actual growth and mutation rates of *L. crispatus* in the human vagina have not been measured, we estimated reasonable *in vivo* doubling times for vaginal *L. crispatus* based on the observed number of SNPs between *L. crispatus* isolates from mother/daughter pair I, the age of daughter I and *L. casei* Zhang mutation rates. Considering the uncertainty in the estimates, transmission of *L. crispatus* from mother to daughter at birth followed by the accumulation of independent mutations during 20 years of persistence in the mother and daughter is a plausible explanation for the observed recombination-filtered SNPs. Future studies comparing genomes of *L. crispatus* isolates from more mother/daughter pairs with a variety of daughter ages are needed.

The 2 *L. crispatus* isolates from the mother had highly similar genomes, differing by only 11 recombination-filtered SNPs. A previous study also observed high similarity between the genomes of multiple vaginal *L. crispatus* isolates from one individual, noting that they were indistinguishable (44). Future investigations of *L. crispatus* genomic variation within an individual may yield further insight on colonization and dynamics of the vaginal microbiota.

Consistent with a previous study, *L. crispatus* isolates from the human vagina were phylogenetically intermixed with isolates from the human urinary tract, including highly similar vaginal (ERS1867668 (SAMEA104208650)) and bladder (ERS1867667 (SAMEA104208649)) isolates from the same subject (Figure 3) (45).

The health implications of vertical transmission of the vaginal microbiota are unknown and were not addressed in this study. However, because vertical transmission seems to be an important factor in determining the composition of the vaginal microbiota there may be important consequences. Vertical transmission of the vaginal microbiota may be one mechanism for maintaining human microbiota over generations via a consistent and specific seeding of the newborn microbiota. Delivery mode is an important factor in determining the early composition of the gut microbiota (46, 47) and is a risk factor for development of immune-related disorders later in life (48). This suggests an important role for the mother’s vaginal microbiota in seeding the infant and setting the stage for development of the gut microbiota. Therefore, maintenance of the vaginal microbiota between generations may be critical for gut microbiota development in each generation.

Additionally, the vaginal microbiota plays an important if not well understood role in reproductive health, with associations between vaginal microbiota composition and infection susceptibility, BV and preterm birth (1–6). Evidence from this study suggests that transmission of microbes from mother to daughter at birth may influence the composition of the daughter’s microbiota later in life and may contribute to the maintenance of specific members of the human vaginal microbiota over generations.

This study provides tantalizing evidence of vertical transmission of the vaginal microbiota. However, this was a small study with only 13 mother/daughter pairs (92% white) and 3/13 daughters born by C-section. Mothers with daughters born by C-section were on average older than mothers with daughters born by vaginal delivery (Table 1, Supplemental Table 1) and two of the three mothers with daughters born by C-section were post-menopausal which could also contribute to a greater difference in mother/daughter vaginal microbiotas (41). Beyond birth mode and reproductive status, other factors including genetics and shared environment could contribute to mother/daughter vaginal microbiota similarity. Of the eleven pairs asked about cohabitation, only one pair (IV) reported that they didn’t currently live together full or part-time (Supplemental Table 1). Therefore, the influence of cohabitation on vaginal microbiota similarity could not be addressed in our study. Genomic analysis of isolates was limited to one member of the vaginal microbiota from 1 mother/daughter pair. Future studies in larger populations, including more racially diverse subjects, more daughters born by C-section and analysis of more isolate genome sequences or metagenomes are required to validate these findings.

## Figures

**Supplemental Figure 1.Principal coordinates analysis (PCoA) of vaginal microbiota from 13 mother/daughter pairs.** The θ_YC_ distances between 101 vaginal microbiota samples are represented by PCoA. Samples from daughters are represented by triangles and samples from mothers by circles. Each mother/daughter pair is represented by a unique color. Biplot arrows represent the 3 OTUs most correlated with position on the PCoA plot.

## List of abbreviations

C-section: Cesarean section
rRNA: ribosomal RNA
OTU: operational taxonomic unit
SNPs: single nucleotide polymorphisms
PCoA: principal coordinates analysis

## Declarations

### Ethics approval and consent to participate

All subjects provided written informed consent. The study was approved by the University of Michigan IRB (HUM00086661).

### Consent for publication

Not applicable.

### Availability of data and material

The raw sequence data generated in this study are available in the NCBI’s SRA:

Bacterial 16S rRNA gene sequences: BioProject PRJNA547595
*L. crispatus* draft genome sequences: BioProject PRJNA547620

GitHub repository:
https://github.com/cbassis/MotherDaughter_Vaginal_Microbiota.study

This repository includes:

- the mothur batch file with steps used to process and analyze 16S rRNA gene sequences
- mothur output files used in final bacterial community analysis and figures
- R code for manuscript figures, statistics and genomic analysis

### Authors’ contributions

CMB was involved in study design and planning, data analysis, figure preparation and manuscript writing. KAB was involved in subject recruitment, sample processing, isolation of *L. crispatus* genomic DNA for sequencing, data analysis and manuscript editing. DES was involved in subject recruitment, sample processing and manuscript editing. KS was involved in genomic data analysis, interpretation of genomic data, phylogenetic tree construction and manuscript editing. AP was involved in genomic data analysis and manuscript editing. ES was involved in genomic data analysis and interpretation and manuscript review. VIA was involved in subject recruitment and planning. EHQ was involved in study design and planning, subject recruitment and manuscript editing. VBY was involved in study design and manuscript editing. JDB was involved with study design and planning, subject recruitment and manuscript editing. All authors read and approved the final manuscript.

## Acknowledgements

We’d like to thank the participants who contributed samples. We’d also like to thank Megan Lemieur for help with study organization and Qualtrics survey implementation and the University of Michigan Microbial Systems Molecular Biology Laboratory for swab DNA isolation, bacterial community and genomic library preparation and sequencing. We’d also like to thank Ingrid Bergin and members of the Young lab for many discussions related to this project over the years. This research was also supported in part through computational resources and services provided by Advanced Research Computing at the University of Michigan, Ann Arbor.

## References

1. Callahan BJ, DiGiulio DB, Goltsman DSA, Sun CL, Costello EK, Jeganathan P, Biggio JR, Wong RJ, Druzin ML, Shaw GM, Stevenson DK, Holmes SP, Relman DA. 2017. Replication and refinement of a vaginal microbial signature of preterm birth in two racially distinct cohorts of US women. Proceedings of the National Academy of Sciences doi:10.1073/pnas.1705899114.

2. Sun S, Serrano MG, Fettweis JM, Basta P, Rosen E, Ludwig K, Sorgen AA, Blakley IC, Wu MC, Dole N, Thorp JM, Siega-Riz AM, Buck GA, Fodor AA, Engel SM. 2022. Race, the Vaginal Microbiome, and Spontaneous Preterm Birth. mSystems 7:e0001722.

3. Klatt NR, Cheu R, Birse K, Zevin AS, Perner M, Noël-Romas L, Grobler A, Westmacott G, Xie IY, Butler J, Mansoor L, McKinnon LR, Passmore J-AS, Abdool Karim Q, Abdool Karim SS, Burgener AD. 2017. Vaginal bacteria modify HIV tenofovir microbicide efficacy in African women. Science (Wash DC) 356:938.

4. McClelland RS, Lingappa JR, Srinivasan S, Kinuthia J, John-Stewart GC, Jaoko W, Richardson BA, Yuhas K, Fiedler TL, Mandaliya KN, Munch MM, Mugo NR, Cohen CR, Baeten JM, Celum C, Overbaugh J, Fredricks DN. 2018. Evaluation of the association between the concentrations of key vaginal bacteria and the increased risk of HIV acquisition in African women from five cohorts: a nested case-control study. The Lancet Infectious Diseases 18:554–564.

5. Bayigga L, Kateete DP, Anderson DJ, Sekikubo M, Nakanjako D. 2019. Diversity of vaginal microbiota in sub-Saharan Africa and its effects on HIV transmission and prevention. Am J Obstet Gynecol 220:155–166.

6. Tamarelle J, Thiébaut ACM, de Barbeyrac B, Bébéar C, Ravel J, Delarocque-Astagneau E. 2019. The vaginal microbiota and its association with human papillomavirus, Chlamydia trachomatis, Neisseria gonorrhoeae and Mycoplasma genitalium infections: a systematic review and meta-analysis. Clinical Microbiology and Infection 25:35–47.

7. The Human Microbiome Project Consortium. 2012. Structure, function and diversity of the healthy human microbiome. Nature 486:207–214.

8. Ravel J, Gajer P, Abdo Z, Schneider GM, Koenig SSK, McCulle SL, Karlebach S, Gorle R, Russell J, Tacket CO, Brotman RM, Davis CC, Ault K, Peralta L, Forney LJ. 2011. Vaginal microbiome of reproductive-age women. Proceedings of the National Academy of Sciences 108:4680–4687.

9. Bassis CM, Allsworth JE, Wahl HN, Sack DE, Young VB, Bell JD. 2017. Effects of intrauterine contraception on the vaginal microbiota. Contraception 96:189–195.

10. O’Hanlon DE, Moench TR, Cone RA. 2013. Vaginal pH and Microbicidal Lactic Acid When Lactobacilli Dominate the Microbiota. PLoS ONE 8:e80074.

11. Tachedjian G, O’Hanlon DE, Ravel J. 2018. The implausible “in vivo” role of hydrogen peroxide as an antimicrobial factor produced by vaginal microbiota. Microbiome 6:29.

12. Dominguez-Bello MG, Costello EK, Contreras M, Magris M, Hidalgo G, Fierer N, Knight R. 2010. Delivery mode shapes the acquisition and structure of the initial microbiota across multiple body habitats in newborns. Proceedings of the National Academy of Sciences 107:11971–11975.

13. Madan JC, Hoen AG, Lundgren SN, et al. 2016. Association of cesarean delivery and formula supplementation with the intestinal microbiome of 6-week-old infants. JAMA Pediatrics 170:212–219.

14. Seekatz AM, Theriot CM, Molloy CT, Wozniak KL, Bergin IL, Young VB. 2015. Fecal Microbiota Transplantation Eliminates *Clostridium difficile* in a Murine Model of Relapsing Disease. Infection and Immunity 83:3838–3846.

15. Kozich JJ, Westcott SL, Baxter NT, Highlander SK, Schloss PD. 2013. Development of a Dual-Index Sequencing Strategy and Curation Pipeline for Analyzing Amplicon Sequence Data on the MiSeq Illumina Sequencing Platform. Appl Environ Microbiol 79:5112–5120.

16. Schloss PD, Westcott SL, Ryabin T, Hall JR, Hartmann M, Hollister EB, Lesniewski RA, Oakley BB, Parks DH, Robinson CJ, Sahl JW, Stres B, Thallinger GG, Van Horn DJ, Weber CF. 2009. Introducing mothur: OpenSource, Platform-Independent, Community-Supported Software for Describing and Comparing Microbial Communities. Appl Environ Microbiol 75:7537–7541.

17. Schloss PD. 2009. A High-Throughput DNA Sequence Aligner for Microbial Ecology Studies. PLoS ONE 4:e8230.

18. Westcott SL, Schloss PD. 2015. De novo clustering methods outperform reference-based methods for assigning 16S rRNA gene sequences to operational taxonomic units. PeerJ 3:e1487.

19. Schloss PD, Westcott SL. 2011. Assessing and Improving Methods Used in Operational Taxonomic Unit-Based Approaches for 16S rRNA Gene Sequence Analysis. Appl Environ Microbiol 77:3219–3226.

20. Wang Q, Garrity GM, Tiedje JM, Cole JR. 2007. Naive Bayesian Classifier for Rapid Assignment of rRNA Sequences into the New Bacterial Taxonomy. Appl Environ Microbiol 73:5261–5267.

21. Cole JR, Wang Q, Fish JA, Chai B, McGarrell DM, Sun Y, Brown CT, Porras-Alfaro A, Kuske CR, Tiedje JM. 2014. Ribosomal Database Project: data and tools for high throughput rRNA analysis. Nucleic Acids Res 42:D633–42.

22. Morgulis A, Coulouris G, Raytselis Y, Madden TL, Agarwala R, Schäffer AA. 2008. Database indexing for production MegaBLAST searches. Bioinformatics (Oxf) 24:1757–1764.

23. Yue JC, Clayton MK. 2005. A similarity measure based on species proportions. Communications in Statistics-Theory and Methods 34:2123–2131.

24. Andrews S. FastQC. http://www.bioinformatics.babraham.ac.uk/projects/fastqc/.

25. Bolger AM, Lohse M, Usadel B. 2014. Trimmomatic: a flexible trimmer for Illumina sequence data. Bioinformatics (Oxf) 30:2114–20.

26. Li H, Durbin R. 2009. Fast and accurate short read alignment with Burrows-Wheeler transform. Bioinformatics (Oxf) 25:1754–60.

27. BWA. http://bio-bwa.sourceforge.net.

28. Picard https://broadinstitute.github.io/picard/.

29. Calling SNPs/INDELs with SAMtools/BCFtools. http://samtools.sourceforge.net/mpileup.shtml.

30. McKenna A, Hanna M, Banks E, Sivachenko A, Cibulskis K, Kernytsky A, Garimella K, Altshuler D, Gabriel S, Daly M, DePristo MA. 2010. The Genome Analysis Toolkit: a MapReduce framework for analyzing next-generation DNA sequencing data. Genome Res 20:1297–303.

31. Pon A, Marcu A, Arndt D, Grant JR, Sajed T, Liang Y, Wishart DS. 2016. PHASTER: a better, faster version of the PHAST phage search tool. Nucleic Acids Res 44:W16–W21.

32. Kurtz S, Phillippy A, Delcher AL, Smoot M, Shumway M, Antonescu C, Salzberg SL. 2004. Versatile and open software for comparing large genomes. Genome Biol 5:R12.

33. Croucher NJ, Page AJ, Connor TR, Delaney AJ, Keane JA, Bentley SD, Parkhill J, Harris SR. 2015. Rapid phylogenetic analysis of large samples of recombinant bacterial whole genome sequences using Gubbins. Nucleic Acids Res 43:e15.

34. Stamatakis A. 2014. RAxML version 8: a tool for phylogenetic analysis and post-analysis of large phylogenies. Bioinformatics (Oxford, England) 30:1312–1313.

35. Wang J, Dong X, Shao Y, Guo H, Pan L, Hui W, Kwok L-Y, Zhang H, Zhang W. 2017. Genome adaptive evolution of Lactobacillus casei under long-term antibiotic selection pressures. BMC Genomics 18:320.

36. Si J, You HJ, Yu J, Sung J, Ko G. 2016. *Prevotella* as a Hub for Vaginal Microbiota under the Influence of Host Genetics and Their Association with Obesity. Cell Host & Microbe 21:97–105.

37. Hickey RJ, Zhou X, Settles ML, Erb J, Malone K, Hansmann MA, Shew ML, Van Der Pol B, Fortenberry JD, Forney LJ. 2015. Vaginal Microbiota of Adolescent Girls Prior to the Onset of Menarche Resemble Those of Reproductive-Age Women. mBio 6:e00097–15.

38. Lee JE, Lee S, Lee H, Song YM, Lee K, Han MJ, Sung J, Ko G. 2013. Association of the vaginal microbiota with human papillomavirus infection in a Korean twin cohort. PLoS One 8:e63514.

39. Kim SJ, Kim SJ, Han K-T, Park E-C. 2017. Medical costs, Cesarean delivery rates, and length of stay in specialty hospitals vs. non-specialty hospitals in South Korea. PLOS ONE 12:e0188612.

40. Caughey AB, Cahill AG, Guise J-M, Rouse DJ. 2014. Safe prevention of the primary cesarean delivery. American Journal of Obstetrics and Gynecology 210:179–193.

41. Brotman RM, Shardell MD, Gajer P, Fadrosh D, Chang K, Silver MI, Viscidi RP, Burke AE, Ravel J, Gravitt PE. 2018. Association between the vaginal microbiota, menopause status, and signs of vulvovaginal atrophy. Menopause 25:1321–1330.

42. Gibson B, Wilson Daniel J, Feil E, Eyre-Walker A. 2018. The distribution of bacterial doubling times in the wild. Proceedings of the Royal Society B: Biological Sciences 285:20180789.

43. Lee YK, Ho PS, Low CS, Arvilommi H, Salminen S. 2004. Permanent Colonization by *Lactobacillus casei* Is Hindered by the Low Rate of Cell Division in Mouse Gut. Appl Environ Microbiol 70:670.

44. Abdelmaksoud AA, Koparde VN, Sheth NU, Serrano MG, Glascock AL, Fettweis JM, Strauss JF, 3rd, Buck GA, Jefferson KK. 2016. Comparison of Lactobacillus crispatus isolates from Lactobacillus-dominated vaginal microbiomes with isolates from microbiomes containing bacterial vaginosis-associated bacteria. Microbiology 162:466–75.

45. Thomas-White K, Forster SC, Kumar N, Van Kuiken M, Putonti C, Stares MD, Hilt EE, Price TK, Wolfe AJ, Lawley TD. 2018. Culturing of female bladder bacteria reveals an interconnected urogenital microbiota. Nat Commun 9:1557.

46. Wampach L, Heintz-Buschart A, Fritz JV, Ramiro-Garcia J, Habier J, Herold M, Narayanasamy S, Kaysen A, Hogan AH, Bindl L, Bottu J, Halder R, Sjöqvist C, May P, Andersson AF, de Beaufort C, Wilmes P. 2018. Birth mode is associated with earliest strain-conferred gut microbiome functions and immunostimulatory potential. Nature Communications 9:5091.

47. Rutayisire E, Huang K, Liu Y, Tao F. 2016. The mode of delivery affects the diversity and colonization pattern of the gut microbiota during the first year of infants’ life: a systematic review. BMC Gastroenterology 16:86.

48. Tamburini S, Shen N, Wu HC, Clemente JC. 2016. The microbiome in early life: implications for health outcomes. Nature Medicine 22:713.

