## Supplementary figures and images for "Vaginal microbiota of adolescents and their mothers: A preliminary study of vertical transmission and persistence"

### Supplemental Figure 1

Supplemental Figure 1

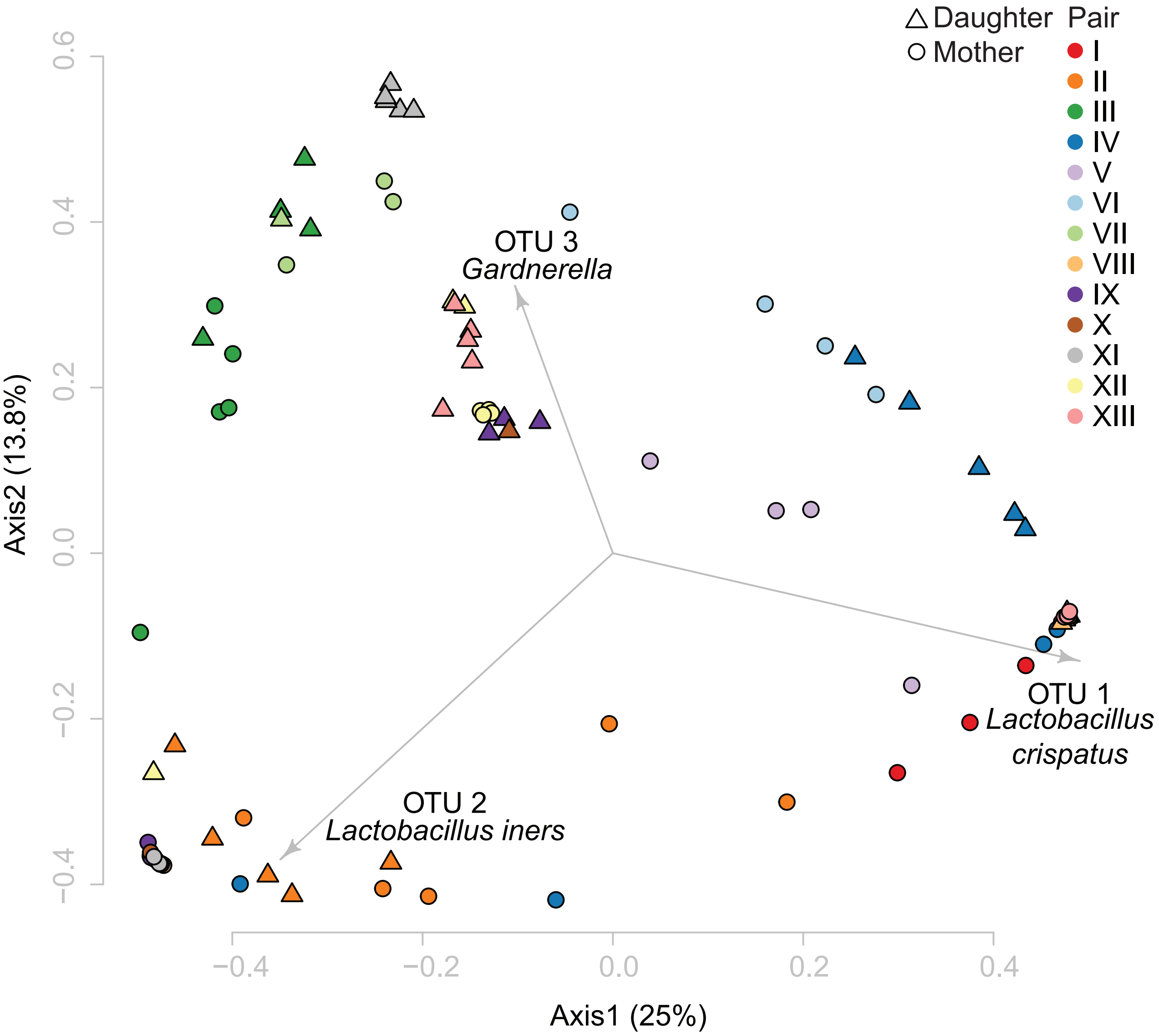
